# Was maize domesticated in the Balsas Basin? Complex patterns of genetic divergence, gene flow and ancestral introgressions among *Zea* subspecies suggest an alternative scenario

**DOI:** 10.1101/239707

**Authors:** Alejandra Moreno-Letelier, J.A. Aguirre-Liguori, Maud I. Tenaillon, Daniel Piñero, Brandon S. Gaut, Alejandra Vázquez-Lobo, Luis E. Eguiarte

## Abstract

The study of maize domestication has overlooked the genetic structure within maize’s wild relative teosinte. Prior to investigating the domestication history of maize (*Zea mays* subspecies *mays*), one should first understand the population history of teosintes and how they relate to maize. To achieve this, we used 32,739 SNPs obtained from a broad sampling of teosinte populations and 46 maize landraces and a) inferred current and past gene flow among teosinte populations and maize, b) analyzed the degree of introgression among *Zea mays* subspecies, and c) explored the putative domestication location of maize. We found geographic structure and introgression between *Zea mays* taxa. Teosinte subspecies have diverged significantly from maize, which indicates that current teosinte populations have evolved mainly independently from maize since the domestication. Our results further suggest that the likely ancestor of maize may maybe have come from Jalisco or the Pacific coast.

**One Sentence Summary:** Shared polymorphism in teosinte suggests a Jalisco origin of maize domestication.

## Introduction

Recent genomic studies on crops and livestock have revealed domestication to be a complex process, where multiple origins and ongoing gene flow are more common than previously thought. For example, gene flow between domesticated and wild forms has been identified in pigs, rice, barley, chili pepper and the common bean, and all of these have been recognized as having multiple centers of domestication (*1*–*4*). In fact, the protracted model of domestication - which proposes that many crops were domesticated over a long period of time with recurrent gene flow- has gained a lot of interest as it can explain the high levels of genetic diversity in many domesticated species, the multiple domestication signatures and ongoing gene flow during a longer transition period (*5*).

*Zea mays* (maize and relatives) consists of three clearly differentiated subspecies: the domestic *Zea mays* spp. *mays* and two wild relatives collectively called teosinte: *Zea mays* spp. *mexicana* (hereafter *mexicana*), and *Zea mays* spp. *parviglumis* (hereafter *parviglumis*). In contrast to complex scenarios proposed for several species, the current consensus on maize domestication points to a single center of origin in the Balsas Basin in Southern Mexico (Fig. 1 populations **1 Tel**, **2 Alch** and **8 Hui**; *6*, *7*; Fig. 1) from the subspecies *parviglumis*. However, inferences of historical distributions from climate data have identified potential refugia from climate change in Western Mexico (Michoacán, Colima and Jalisco) during the warm period of the mid-Holocene which coincides with the inferred dates of maize domestication (*8*).

**Fig. 1.**
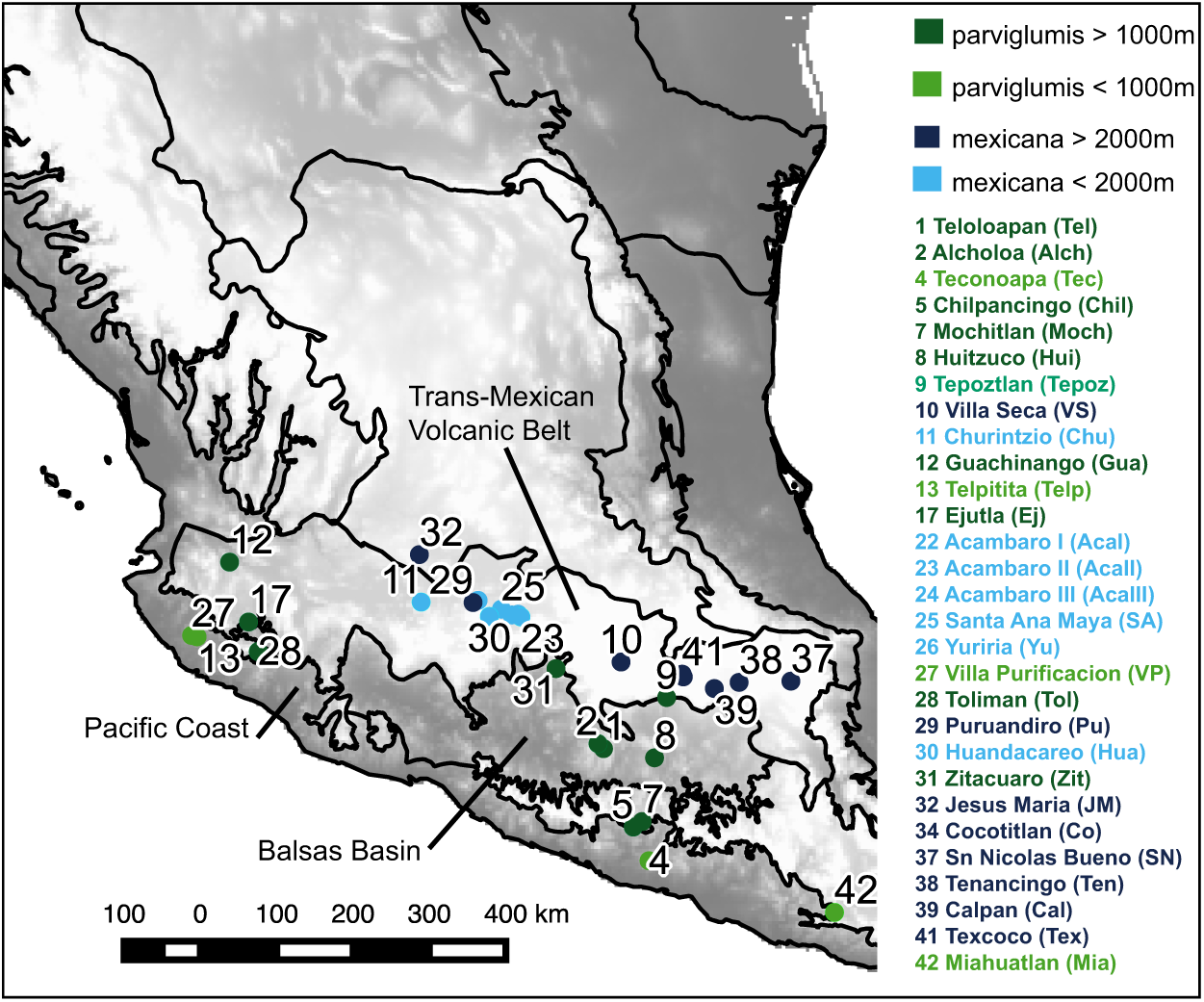
Distribution of sampling sites of both analyzed teosinte varieties. The main biogeographic regions represented in this study are highlighted. Numbers indicate the different populations, with populations names and abbreviations on the right. The subspecies and the altitudinal distribution are indicated by shades of green and blue.

Additionally, this area has been identified as the center of domestication of two species of bean (*Phaseolus vulgaris* and *P. acutifolius*; *4*, *9*, *10*), and archeological evidence of early agriculture has been found for the Early-Holocene (*11*). The domestication in the Balsas Basin has been supported by genetic analyses (*6*), however, those studies have included a limited sampling of *parviglumis* populations from Western Mexico (Jalisco, Michoacán, Colima). Genetic studies that have included Western populations have uncovered a high level of genetic variation and strong genetic differentiation in these populations (*12*, *13*). Moreover, van Heerwarden et al. (*7*) suggested ancestral teosinte alleles in the Western region, rather than in the Balsas Basin.

Here we explored alternative scenarios of maize domestication (Balsas vs. Jalisco-Pacific) and introgression between wild and cultivated forms by including 29 teosinte populations from across the entire range of *parviglumis* and *mexicana,* as well as 46 maize landraces. We first assessed genetic structuring of teosinte populations using 32,739 Single Nucleotide Polymorphisms (SNPs), and inferred migration events using a maximum-likelihood framework (*12*), with *Tripsacum dactyloides* as an outgroup to infer ancestral states of alleles and distinguish gene flow from shared ancestral polymorphism (*14*).

## Results

### Genetic differentiation and genetic structure

We assessed genetic structuring representing 403 samples from 28 teosinte populations, and 46 maize landraces (*15*) using a Principal Component Analysis of 32 739 SNPs. This analysis revealed several interesting features (Fig. 2). First, there is a clear differentiation of the Mexican maize landraces from their wild progenitor. Second, there is high clustering and low level of genetic differentiation among *mexicana* populations. Third, in contrast to *mexicana*, *parviglumis* populations are more differentiated, with the Jalisco populations being a prime example, particularly populations **13Telp** and **27VP** (Fig. 1;*12*, *13*). The dendrogram constructed using Nei’s genetic distance indicates that individuals are strongly clustered by population (Fig. 3), and that the Guachinango population (**12Gua**) is early divergent and closer to maize.

**Fig.2.**
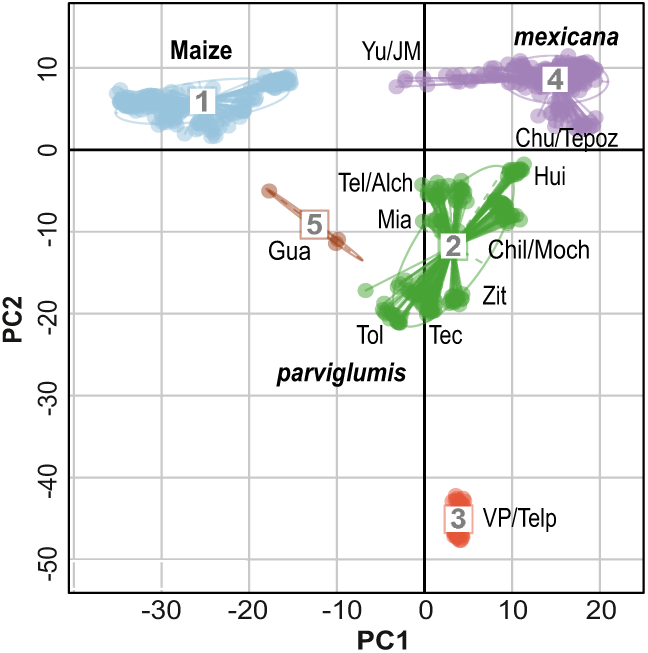
Principal component analysis of 32739 SNPs. Clusters were constructed using UPGMA based on Euclidean distances of the first two components. The populations are identified by the abbreviations used in Fig.1.

**Fig. 3.**
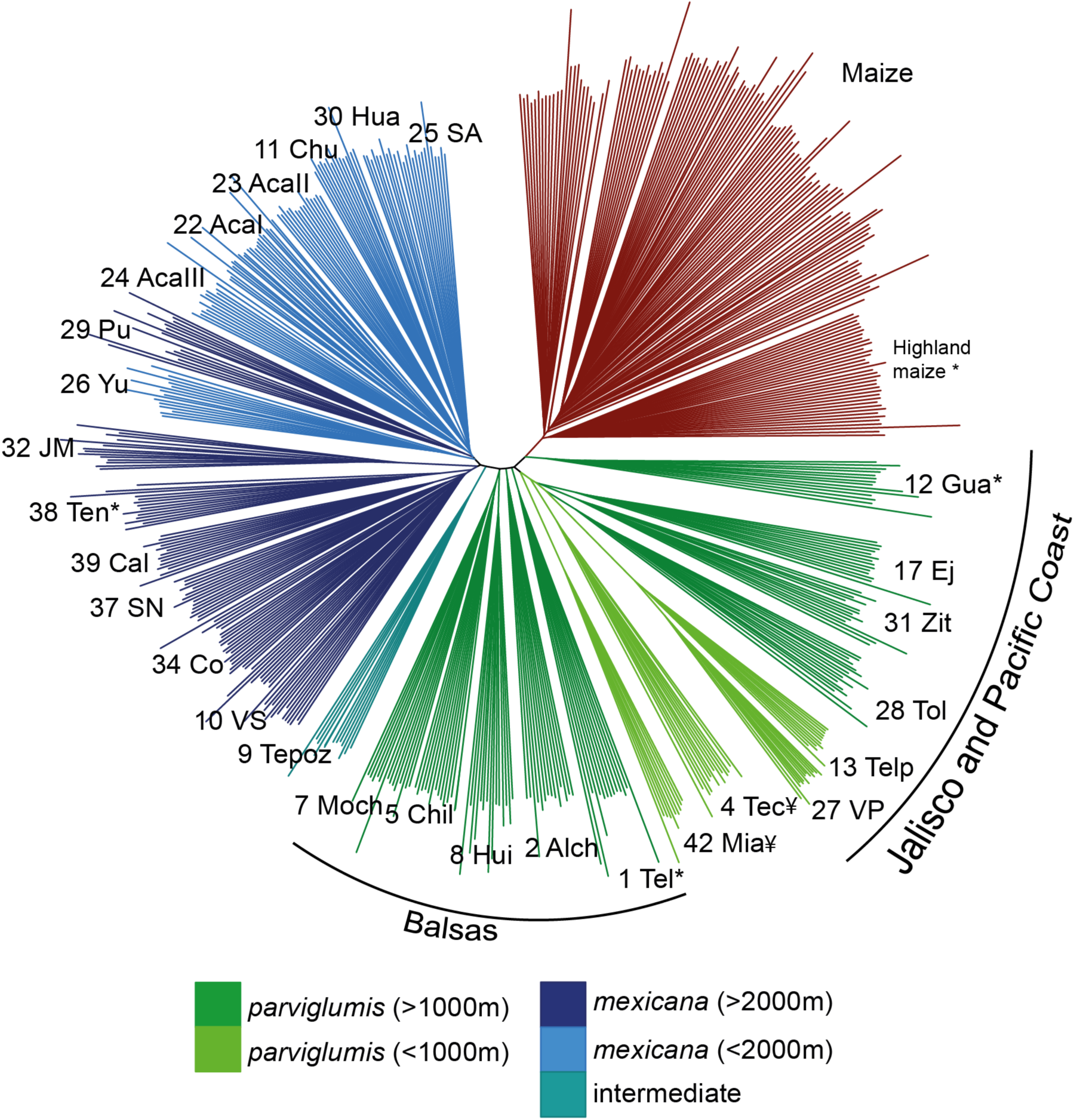
Dendrogram of Nei’s genetic distance of maize and teosinte samples. The colors denote the variety and altitudinal distribution of teosinte varieties. Population with asterisks were used to infer ancestral introgression in other tests. 
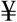
 Southern Guerrero and Oaxaca populations.

Genetic differentiation between teosinte subspecies and maize was also apparent in the genetic clustering analysis implemented by fastStructure where *K*=3 was the optimal value for the number of clusters, followed by *K*=4. For *K*=3, maize and *mexicana* appear fairly homogeneous, while *parviglumis* shows a high degree of admixture in some populations, with the exception of populations **13Telp** and **27VP**, which correspond to the Jalisco populations of Telpitita and Villa Purificacion (Fig. S1). For *K*=4, further population structure is found in *parviglumis*, which coincides with the results seen in the PCA. Overall, genetic diversity analyses tend to indicate that populations from Jalisco group more closely with maize; fastStructure suggests that some but not all of these populations may be admixed with maize.

### Gene flow and shared ancestral polymorphism

To further investigate the possibility of introgression, we employed TreeMix analyses, using *Tripsacum* as an outgroup. The use of *Tripsacum* instead of other members of the genus *Zea* was important to ensure no contemporary gene flow could produce artifacts in the inference of migration events and shared ancestral polymorphism. As no significant genetic structure was detected within maize landraces (*15*), we used all 161 individuals from 46 landraces as one population for all analyses. In the population graph, *mexicana* populations appear nested within *parviglumis*, with the Jalisco populations of *parviglumis* being ancestral (**12Gua** and **28Tol**; Fig. 4 and Table S1). Ancestral polymorphism is shown as a higher than expected covariance of allele frequencies between *Tripsacum* and maize and *parviglumis* populations in the first column of the covariance matrix estimated by Treemix (Fig. 4, cooler colors). Once ancestral polymorphism was identified, two potential admixture events were then detected (arrows in Fig. 4), one from a highland *mexicana* population (**32JM**, Jalisco) to maize and another from maize to another *mexicana* highland population (**38Ten**, Tlaxcala).

**Fig. 4.**
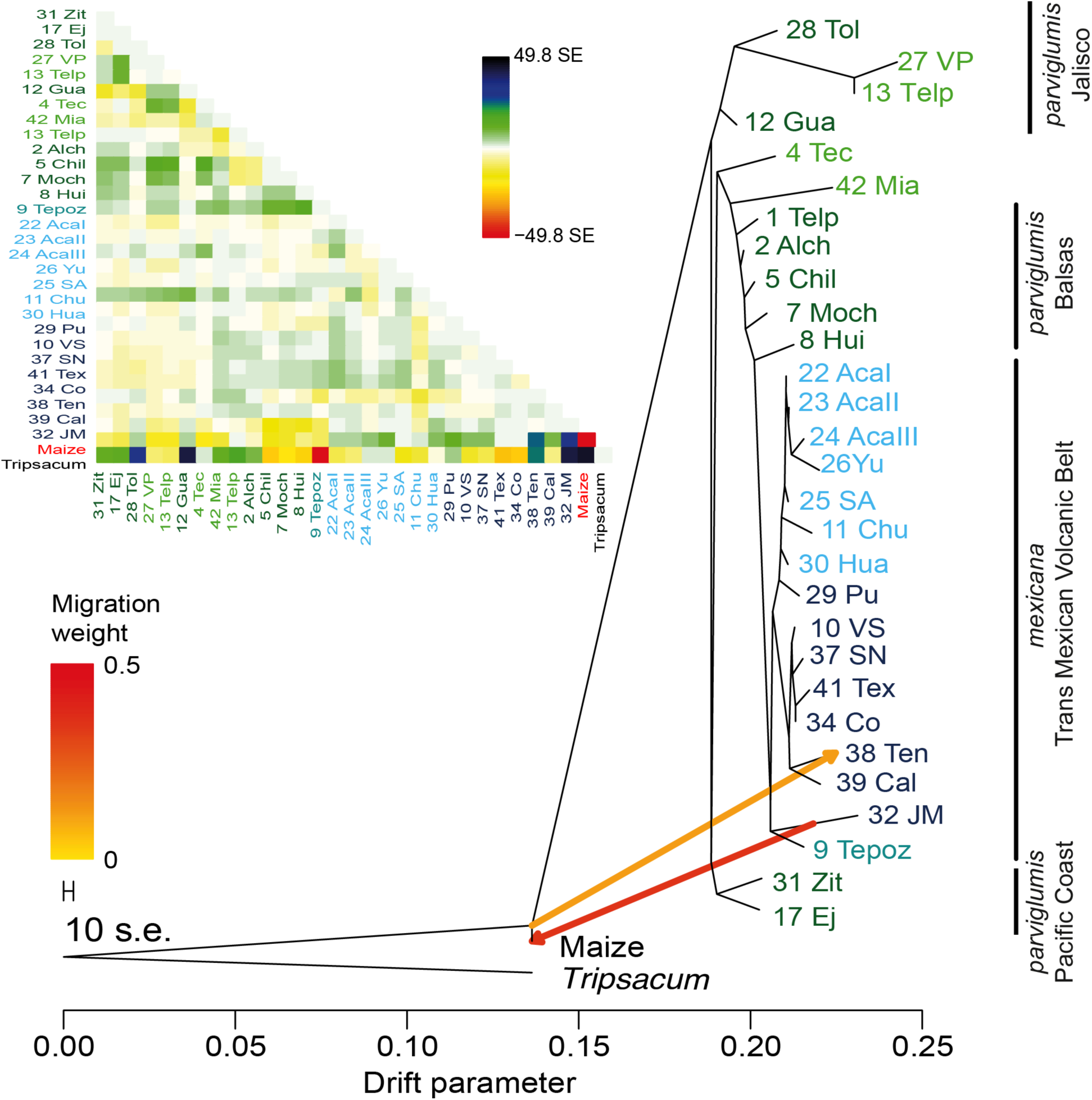
Population graph and covariance matrix obtained with TreeMix with 30673 SNPs. Two migration events were inferred once shared polymorphism with *Tripsacum* was considered, and can be seen in blue-black colors in the covariance matrix. The drift parameter indicates the intensity of genetic drift. The color of the migration lines indicates the percentage of loci shared between populations. The migration weight from maize to **38Ten** is 0.096 (9.6%) and between **12JM** and maize is 0.47 (47%). Populations with asterisks were used to infer ancestral introgression in other tests.

An additional introgression analysis (ABBA-BABA; *17*) was performed to test for geneflow between *parviglumis* from Jalisco populations (**12Gua**), Balsas (**1Tel**), a highland *mexicana* population (**38Ten**) and highland maize. The *D-statistic* detects deviations from random lineage sorting and *F*_*d*_ which estimates the proportion of introgressed alleles (*17*). In all cases, *Tripsacum* was used as an outgroup and of the 24 possible 4 taxa combinations, only two showed significant deviations from the null ABBA-BABA distribution. The first four taxon tree (((**1Tel**,**12Gua**)maize)*Tripsacum*) had a significant *D-statistic* (*p*=0.0335) between **12Gua** and maize, and a *F*_*d*_ value that corresponds to 10% of shared alleles (Fig. 5a). The second tree (((**1Tel**,**38Ten**)maize)*Tripsacum*) had a non-significat *D-statistic* (*p*=0.1) but a significant *F*_*d*_ value (Z-score=2.412) corresponding to 16% shared alleles between **38Ten** and maize (Fig. 5b), similar to what was obtained with Treemix. No four-taxon combinations recovered significant signals of introgression between Balsas (**1Tel**) and maize.

**Fig. 5.**
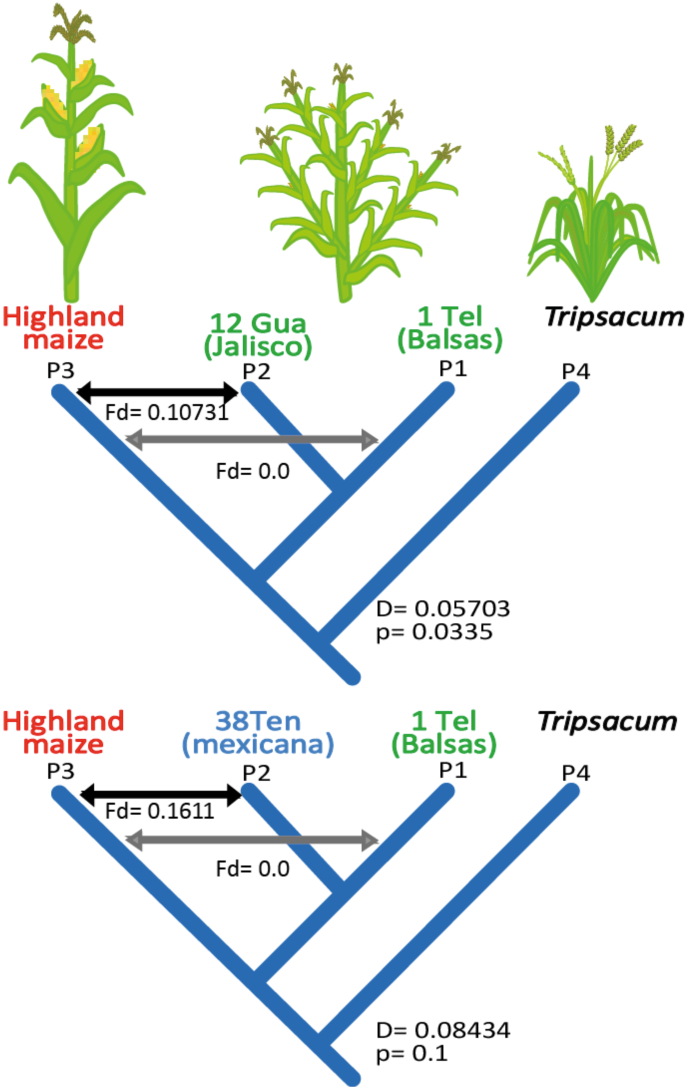
Four taxon ABBA-BABA introgression tests that yielded significant results. a) Significant positive *D* value means introgression between highland maize and **12 Gua** (Jalisco), *F_d_* denotes de proportion of shared alleles. b) Non-significant *D* value between Highland maize and **38 Ten**, but significant *F*_*d*_ value (*Z-score*= 2.4) equating to 16% of shared alleles.

## Discussion

The PCA results show a high degree of genetic structure and differentiation of *parviglumis* populations (Fig. 2). Particularly the Jalisco lowland populations of Telpitita (**13Telp**) and Villa Purificación (**27VP**) on the Pacific Coast form a distinct cluster. This genetic heterogeneity highlights the importance of having a dense enough sampling in the mountains of Jalisco and neighboring Michoacán, which were not considered in previous studies as a potential center of domestication, despite the area’s high biological and cultural diversity (*11*, *12*). In contrast to *parviglumis*, maize and *mexicana* form relatively homogeneous clusters in the PCA respectively (Fig. 2, clusters 1 and 4). The clear genetic differentiation between maize and teosinte is also found in the fastSTRUCTURE analysis (Fig. S1).

Despite the differentiation within teosinte, *mexicana* is nested within *parviglumis*, and both teosinte subspecies are genetically differentiated from maize (Figs. 3-4). These results are different from studies conducted using microsatellites and a smaller teosinte sample, where maize was nested within *parviglumis* and *Zea mays* spp. *huehuetenangensis* was used as an outgroup (*6*). This difference in topologies could be due to different factors: 1) The inclusion of Jalisco populations here, which have not been used previously in other studies; 2) Extreme genetic drift and isolation in *huehuetenangensis* resulting in many private alleles and genetic divergence. These analyses also highlight the basal position of Jalisco-Pacific populations with respect to the Balsas populations from Guerrero. In order to evaluate the utility of using other members of the *Zea* genus as outgroups, we performed a Principal Component Analysis and a constructed a Nei’s genetic distance dendrogram with GBS data published by Swarts et al. (*18*). Our results show that despite *Z. luxurians* and *Z. mays* subsp. *huetentenanguensis* forming distinctive clusters in the PCA (Fig. S2), they group with *parviglumis* (Fig. S3), and are not reliable outgroups. Furthermore, there is evidence of natural hybridization between all diploid members of *Zea* (*19*), so using *Tripsacum* as an outgroup is a better option to correctly infer the relationships within teosinte. Overall, the GBS dendrogram (Fig. S3) confirms our results that maize has diverged significantly from teosinte, instead of being nested within it, a result also confirms the results of Buckler and Edward with ITS markers (*20*)

The Treemix analysis detected only two migration events that showed marked deviations to the expected population covariance between maize and *mexicana* (Fig. 4). Introgression between maize and teosinte has been widely reported, but previous studies could not differentiate between contemporary processes and ancestral introgression (*7*, *21*). Our migration inference with Treemix identifies two ancestral migration events in opposite directions: one from ancestral maize to a *mexicana* population in Tlaxcala (**38 Ten**), and another from an ancestral *mexicana* from Jalisco to maize. The ancestral nature of these events is determined by the position of the migration arrow along the branch, not near the tip of the modern populations, and has been corroborated by other studies (*7*, *14*, *21*). Currently, introgression happens from teosinte to maize, however, the inferred weight of the migration from the ancestral maize into *mexicana* was 9.6%, far higher than the 1-2% obtained experimentally (Fig. 4; *19*). Therefore, introgression probably mainly occurred at a time when reproductive incompatibilities between maize and teosinte were not yet as developed. Likewise, current estimates of gene flow from *mexicana* to maize range from 4-20% (*21*), yet we detected a migration weight of 47% between one *mexicana* population (**32JM**) from Jalisco and maize. Overall, there are more introgressed alleles from *mexicana* to maize, than the opposite direction.

Introgression between *mexicana* and maize has been proposed to be the source of adaptive alleles to survive in highland environments (*21*). However, by including *Tripsacum* in this study we could detect shared ancestral polymorphism and pinpoint the source and scope of that introgression. The high migration weight detected in our study suggests that the contribution of *mexicana* to the contemporary landraces of maize has been substantial, even if the initial domestication was from *parviglumis*.

A recent study comparing the genomes of maize specimens from ~5000 yr B.P with modern maize and teosinte, found that domestication was not yet complete and possibly there was no reproductive isolation between maize and teosinte (*22*). Ongoing gene flow during domestication has been widely documented in animals (*23*, *24*), and it may account for 10% of the genome of cultivated sunflower (*25*). This introgression can only occur in areas where the domesticated forms coexist with their wild relatives, and requires ongoing selection to preserve desired traits, as it is the case of maize and teosinte (*21*).

Finally, our results from the ABBA-BABA test show that *parviglumis* from Jalisco-Pacific has introgressed with maize, while the Balsas population shows no such pattern (Fig. 5a), as it has been usually proposed (*6*). Introgression between *parviglumis* and maize is not recovered by Treemix, but it is important to highlight that Treemix is based on allele frequency covariance, estimated using the entire sample of maize. For the ABBA-BABA test, only highland maize landraces produced significant results, which have been proposed to be the first to have been domesticated (*26*). When performing the same tests with a random sample of all landraces (not only highland), no tests were significant. Furthermore, it is more likely to assume that the introgression between **12Gua** and highland maize is ancient, given the ecological divergence of modern highland maize from the Central Plateau and lowland *parviglumis*. No introgression was detected between **12Gua** and lowland and midland maize. Therefore, this introgression must have happened before maize introgressed with highland *mexicana* and was cultivated in Central Mexico highlands.

Our result is different from previous studies that did not sample the Jalisco region extensively, taking into account that not all teosinte populations might have served as genetic sources of maize. Unlike findings from studies with smaller geographical samples (*6*, *22*), maize is not nested within teosinte, and it clearly forms a separate clade (Figs. 3 and S3). This result suggests that modern teosinte continued its independent evolution after the initial domestication. To prove this hypothesis, it would be necessary to gain access to ancient teosinte to correctly place the maize lineage within the teosinte genealogy including the archeological samples from Western Mexico (*11*).

Some authors have regarded Jalisco as an important center of domestication in Mesoamerica because the wild relatives of all main crops (maize, squash and beans) are distributed there and at least one of them (beans) were domesticated in the region (*11*). Moreover, genetic analyses of *P. vulgaris* show that the Jalisco accessions are basal to cultivated bean (*9*, *10*, *27*), just as the *parviglumis* population of Guachinango (**12Gua**) is early divergent in our dendrogram and population graph (Figs. 2 and 3), and it is early divergent in the re-analized GBS data (Fig. S3). Unfortunately, the inherent ascertainment bias of our data prevents us from estimating accurate divergence dates (*28*), however, a whole genome genotyping approach together with ancient DNA techniques could help us finally answer the question of the time and place of maize domestication.

Our results reveal an ancient divergence between maize and teosinte, with Jalisco-Pacific populations being early divergent, and Balsas populations derived. This pattern suggests that a population from Western Mexico may be a better candidate for the center of maize domestication than the Balsas basin, as this region is more diverse and shares more ancestral polymorphism with maize and close relative *Tripsacum dactiloydes*. In addition, we found evidence for intense ancient gene flow between maize and *mexicana*: first, from a *mexicana* highland population from Jalisco to maize, which has likely contributed to maize adaptation to Mexican central highlands; second, from maize to a *mexicana* highland population of the Trans Mexican Volcanic Belt. Again, by incorporating a more comprehensive sampling of *mexicana* populations, we obtained higher estimates of gene flow than in previous studies (*21*). Such gene flow has been the likely cause of promoted diversification of maize landraces.

## Acknowledgments

This work was supported by grant CB2011/167826 (CONACYT Investigación Científica Básica) to LEE, grant CN-10-393 (UC MEXUS-CONACYT) to LEE and BSG, and grant M12-A03 ECOS Nord France -CONACYT-ANUIES 207571 to LEE and MIT. Maize landraces collection and genotyping was funded by Secretaría de Medio Ambiente y Recursos Naturales (SEMARNAT) through a grant to CONABIO awarded to DP. We are grateful to Erika Aguirre-Planter and Laura Espinosa-Asuar for technical support, and to Jeffrey Ross-Ibarra for his comments that helped improve this manuscript. This paper was written during a sabbatical leave of LEE in the University of Minnesota, Department of Plant and Microbial Biology in Peter Tiffin laboratory, with support by scholarships from PASPA, DGAPA, UNAM. All teosinte and *Tripsacum dactyloides* genotypes are available in the Dryad Digital repository (http://datadryad.org/resource/XXX), and maize landrace genotypes are available in http://datadryad.org/resource/doi:10.5061/dryad.4t20n.

## Supplementary Materials

### Materials and Methods

#### Sampling

Plant material was collected from 28 populations of teosinte, 15 populations of *mexicana* and 14 populations of *parviglumis* for a total of 403 individuals, which covers the entire range of geographical and environmental conditions of both subspecies. Details of the sampling design, DNA extraction, genotyping and locality information are described in (*13*, *29*), and in the Supplementary Material.

Maize samples comprised 161 individuals from 46 landraces from all over Mexico, averaging 3.6 individuals per landrace. Information about sampling, DNA extraction and genotyping can be found in (*15*).

#### SNP calling and data quality assessment

Single nucleotide polymorphisms were detected with the Illumina Maize SNP50 Bead Chip as described for maize by Arteaga et al. (*15*) and for teosinte in Aguirre-Liguori et al. (*13*). Automated allele calling was performed using GenomeStudio 2010.1 (Genotyping module 1.7.4; Illumina), excluding those loci with a GenTrain score <0.3 and those with more than 10% of missing data. Scores between 0.3-0.45 were manually checked and curated. After those filters, a total of 34,981 SNPs were recovered. To reduce the redundancy of the data, loci in strong linkage disequilibrium (r^2^>0.8) were removed using Plink 1.07 (*30*), for a total remaining of 32,739 loci.

#### Genetic differentiation and genetic structure

Genetic differentiation was explored with a Principal Component Analysis with all the 32,739 loci of maize and teosinte, using the package Adegenet and ade4 (*31*) implemented in R (*32*). The clustering analysis was performed with UPGMA implemented by ade4 (*33*). Nei’s genetic distances between individuals were computed using the R package *ape*, and used to construct a Neighbor-Joining dendogram with the same package (*34*).

Genetic clustering analyses were performed with fastSTRUCTURE (*35*). The number of *K* values tested ranged from 1 to 10, with 10 iterations. The optimal *K* values were evaluated with the Evanno method (*36*)

#### Gene flow and ancestral introgression

It is difficult to discern between shared ancestral polymorphism and gene flow (*37*). To do so, we used Treemix, which is based on a maximum-likelihood population tree and identifies pairs of populations with a higher covariance of allelic frequencies than expected under the no migration model (*14*), and determines ancestral allele frequencies by using a suitable out-group, *Tripsacum dactyloides* (data from the 50K SNP dataset from (*12*)). The analysis was carried out with a dataset of 30,673 SNPs that were shared between teosinte, maize and *T. dactyloides*.

To further explore gene flow between populations, especially with highland maize landraces, we performed an ABBA-BABA test which estimates de *D*-statistic based on the distribution of ABBA and BABA polymorphisms in a four taxa tree with *T. dactyloides* as an outgroup. The *D*-statistic estimates the deviation from the expected random distribution of ABBA-BABA polymorphism in a four taxon tree. We also estimated the proportion of introgressed loci using the *f*_*d*_ statistic (*17*). We tested two introgression hypotheses: 1) Balsas (**1 Teloloapan**) and Jalisco (**12 Guachinango**) with maize; 2) Balsas (**1 Teloloapan**) and highland *mexicana* (**38 Tenancingo**) withmaize. The teosinte samples included 10 individuals per population, and the maize sample was performed using 10 individuals from highland landraces,which were early divergent in our distance analyses. A second analysis was performed with a random sample of 40 individuals from all landraces (lowland, midland and highland), but those tests were not significant. All non-variable loci in the sample were removed, yielding a total of 26,162 loci and were analyzed in blocks of 1000 using the package HybridCheck (*16*).

#### GBS data analyses

The analyses of GBS teosinte and maize data from Swarts et al. (*18*) were performed by filtering all sites with over 50% of missing data using and those with missing allele frequencies under 0.05 using vcftools (http://vcftools.sourceforge.net/). That yielded a total of 165 192 SNPs, but after removing sites with gaps, the total was 122 085 SNPs. The Principal Component analysis was done with the glPca function of the Adegenet package (*31*), forcing the number of alleles to be centered and scaled. We retained 10 principal components that represented 17.9% of the total variance. Finally, we plotted the first two principal components, assigning to each dot the race or subspecies to which they belong. The Neighbor-joining dendrogram was constructed from a Nei’s distance matrix obtained from the PCA results described before, using the ape package in R (*34*).

**Author contributions**:AML performed the analyses and wrote the manuscript; LEE, BSG and MIT designed the study and improved the manuscript; JAL and AVL performed lab work and preliminary bioinformatics analyses; DP provided all the maize samples and SNP data.

**Fig. S1.**
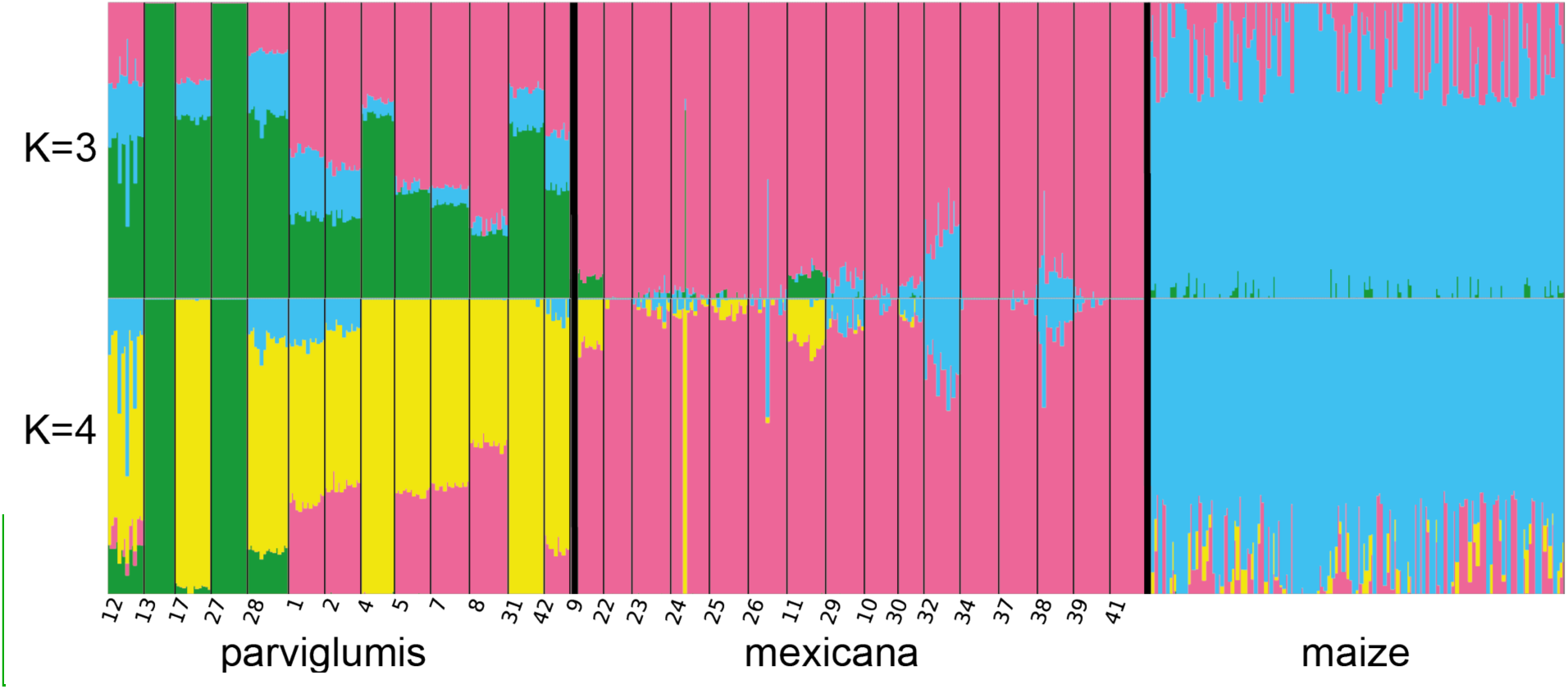
FastStructure analysis carried out with the full dataset. Best *K* values were *K*=3 and *K*=4.

**Fig. S2.**
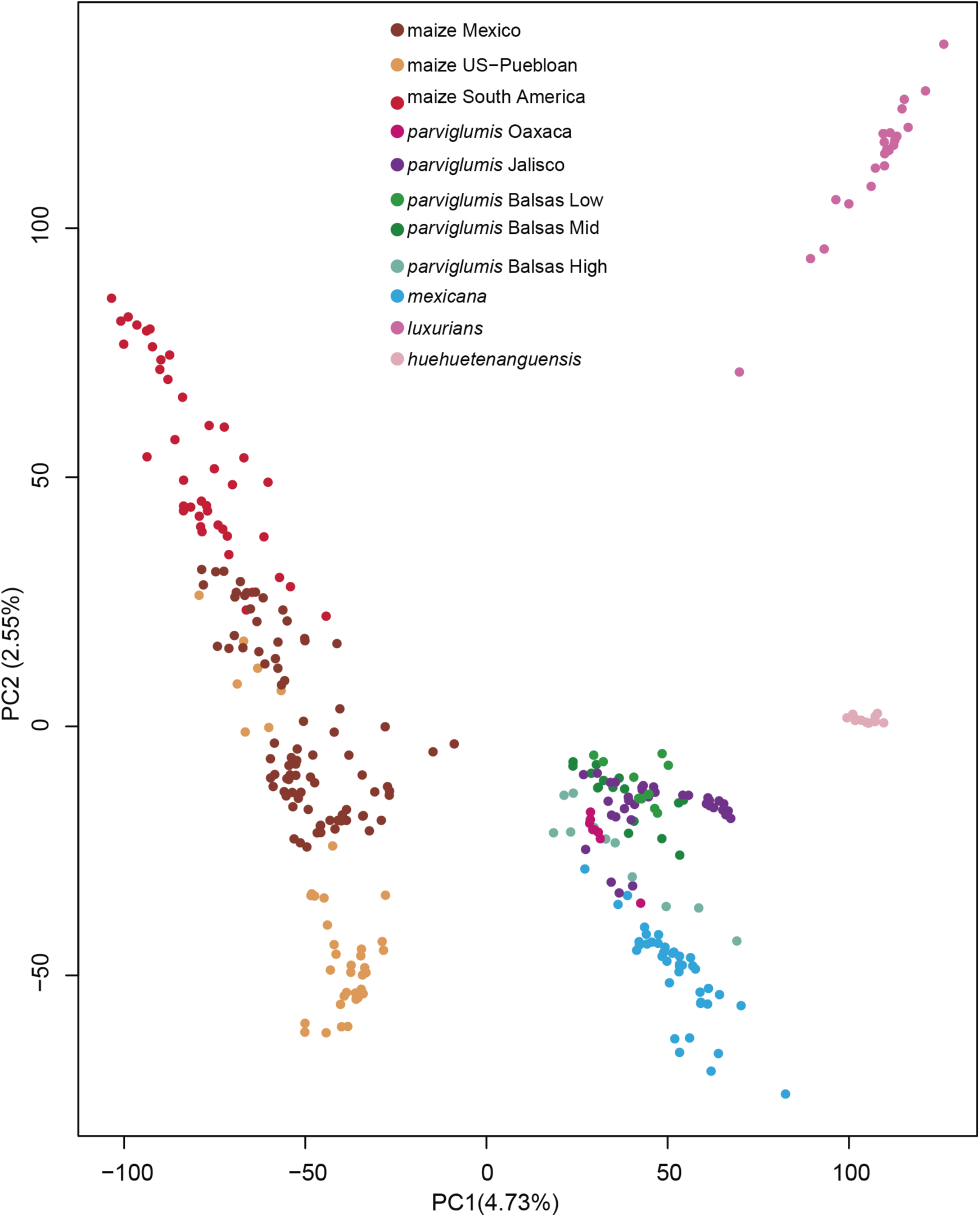
Principal component analysis of 12, 085 SNPs originally published by Swarts et al. (2017)

**Fig. S3.**
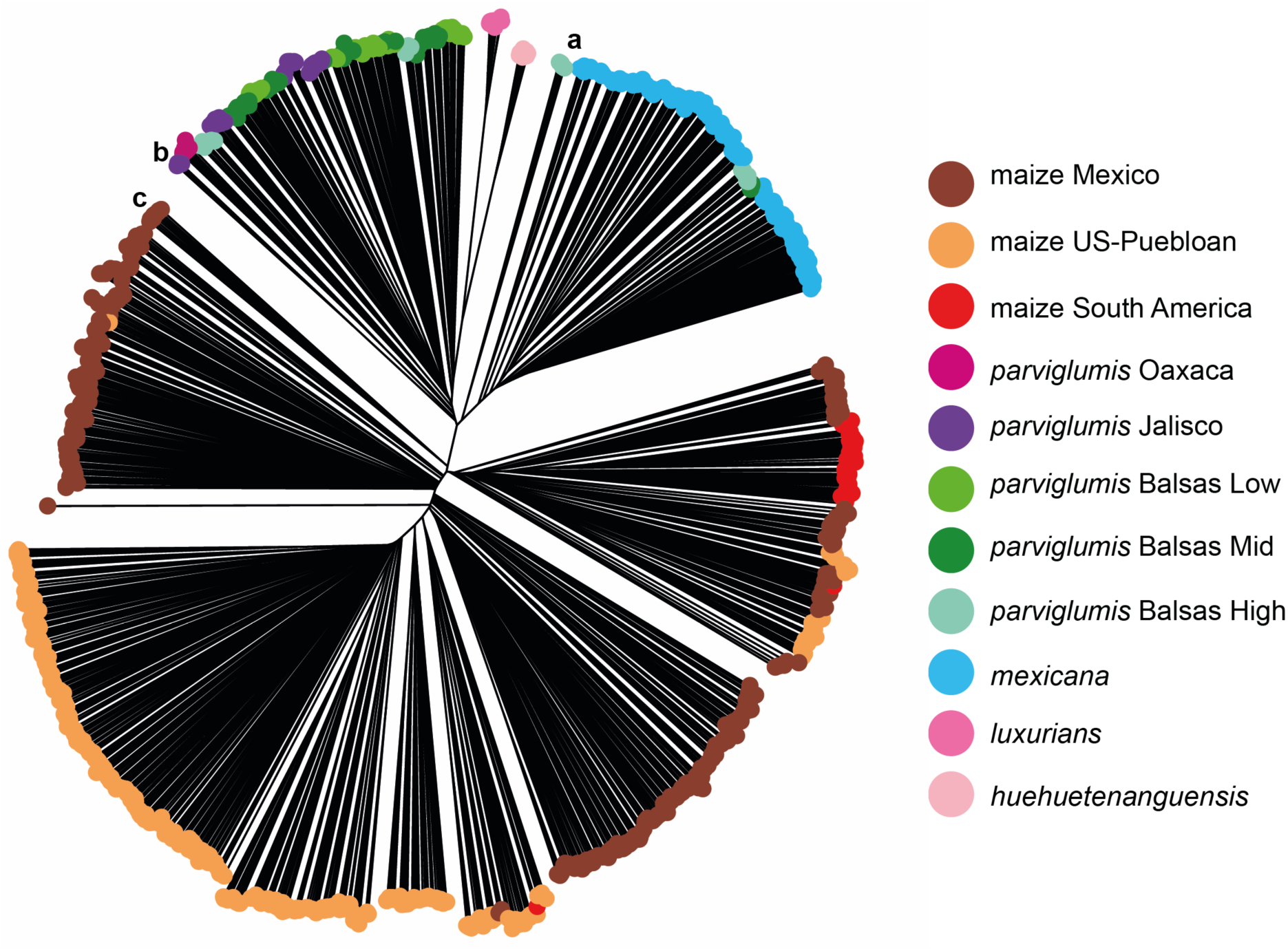
Neighbor-joining dendrogram of 122 085 SNPs from Swarts et al. (*18*). Note the close relationship of *Zealuxurians*, *Zea mays* subsp. *huehuetenanguensis* and *Zea mays* subsp. *parviglumis.* The labels correspond sosampling sites close to those used in our study: a) Site 15km North of **1Teloloapan**; b) site 16 km West of **12Guachinango**; c) Highland maize landraces.

## References and Notes

G. Larson et al., Worldwide Phylogeography of Wild Boar Reveals Multiple Centers of Pig Domestication. Science. 307, 1618–1621 (2005).

B. L. Gross, Z. Zhao, Archaeological and genetic insights into the origins of domesticated rice. Proc. Natl. Acad. Sci. U. S. A. 111, 6190–7 (2014).

K. H. Kraft et al., Multiple lines of evidence for the origin of domesticated chili pepper, Capsicum annuum, in Mexico. Proc. Natl. Acad. Sci. U. S. A. 111, 6165–6170 (2014).

E. Bitocchi et al., Molecular analysis of the parallel domestication of the common bean (Phaseolus vulgaris) in Mesoamerica and the Andes. New Phytol. 197, 300–13 (2013).

R. G. Allaby, D. Q. Fuller, T. a Brown, The genetic expectations of a protracted model for the origins of domesticated crops. Proc. Natl. Acad. Sci. U. S. A. 105, 13982–13986 (2008).

Y. Matsuoka et al., A single domestication for maize shown by multilocus microsatellite genotyping. Proc. Natl. Acad. Sci. U. S. A. 99, 6080–6084 (2002).

J. van Heerwaarden et al., Genetic signals of origin, spread, and introgression in a large sample of maize landraces. Proc. Natl. Acad. Sci. U. S. A. 108, 1088–1092 (2011).

M. B. Hufford, E. Martínez-Meyer, B. S. Gaut, L. E. Eguiarte, M. I. Tenaillon, Inferences from the historical distribution of wild and domesticated maize provide ecological and evolutionary insight. PLoS One. 7, e47659 (2012).

M. Kwak, J. A. Kami, P. Gepts, The putative Mesoamerican domestication center of Phaseolus vulgaris is located in the Lerma – Santiago Basin of Mexico. Crop Sci. 49, 554–563 (2009).

R. Papa, P. Gepts, Asymmetry of gene flow and differential geographical structure of molecular diversity in wild and domesticated common bean (Phaseolus vulgaris L.) from Mesoamerica. Theor. Appl. Genet. 106, 239–250 (2003).

D. Zizumbo-Villarreal, P. Colunga-GarcíaMarín, Origin of agriculture and plant domestication in West Mesoamerica. Genet. Resour. Crop Evol. 57, 813–825 (2010).

T. Pyhäjärvi, M. B. Hufford, S. Mezmouk, J. Ross-Ibarra, Complex patterns of local adaptation in teosinte. Genome Biol. Evol. 5, 1594–1609 (2013).

J. Aguirre-Liguori et al., Connecting genomic patterns of local adaptation and niche suitability in teosintes. Mol. Ecol. 26, 4226–4240 (2017).

J. K. Pickrell, J. K. Pritchard, Inference of population splits and mixtures from genome-wide allele frequency data. PLoS Genet. 8, e1002967 (2012).

M. C. Arteaga et al., Genomic variation in recently collected maize landraces from Mexico. Genomics Data. 7, 38–45 (2016).

B. J. Ward, C. van Oosterhout, Hybridcheck: Software for the rapid detection, visualization and dating of recombinant regions in genome sequence data. Mol. Ecol. Resour. 16, 534–539 (2016).

S. H. Martin, J. W. Davey, C. D. Jiggins, Evaluating the use of ABBA-BABA statistics to locate introgressed loci. Mol. Biol. Evol. 32, 244–257 (2015).

K. Swarts et al., Genomic estimation of complex traits reveals ancient maize adaptation to temperate North America. 515, 512–515 (2017).

M. B. Hufford, P. Bilinski, T. Pyhäjärvi, J. Ross-Ibarra, Teosinte as a model system for population and ecological genomics. Trends Genet. 28, 606–615 (2012).

E. S. Buckler IV, T. P. Holtsford, Zea systematics: Ribosomal ITS evidence. Mol. Biol. Evol. 13, 612–622 (1996).

M. B. Hufford et al., The genomic signature of crop-wild introgression in maize. PLoS Genet. 9, e1003477 (2013).

M. Vallebueno-Estrada et al., The earliest maize from San Marcos Tehuacán is a partial domesticate with genomic evidence of inbreeding. Proc. Natl. Acad. Sci. U. S. A. 113, 14151–14156 (2016).

L. A. F. Frantz et al., Evidence of long-term gene flow and selection during domestication from analyses of Eurasian wild and domestic pig genomes. Nat. Genet. 47, 1141–1148 (2015).

S. D. E. Park et al., Genome sequencing of the extinct Eurasian wild aurochs, Bos primigenius, illuminates the phylogeography and evolution of cattle. Genome Biol. 16, 234 (2015).

G. J. Baute, N. C. Kane, C. J. Grassa, Z. Lai, L. H. Rieseberg, Genome scans reveal candidate domestication and improvement genes in cultivated sunflower, as well as post-domestication introgression with wild relatives. New Phytol. 206, 830–838 (2015).

F. Freitas, G. Bendel, R. Allaby, T. Brown, DNA from primitive maize landraces and archaeological remains: implications for the domestication of maize and its expansion into South America. J. Achaeological Sci. 30, 901–908 (2003).

M. Chacón, B. Pickersgill, D. Debouck, J. Arias, Phylogeographic analysis of the chloroplast DNA variation in wild common bean (Phaseolus vulgaris L.) in the Americas ´. Plant Syst. Evol. 266, 175–195 (2007).

A. Keinan, J. C. Mullikin, N. Patterson, D. Reich, Measurement of the human allele frequency spectrum demonstrates greater genetic drift in East Asians than in Europeans. Nat. Genet. 39, 1251–1255 (2007).

C. M. Diez et al., Genome size variation in wild and cultivated maize along altitudinal gradients. New Phytol. 199, 264–276 (2013).

S. Purcell et al., PLINK: a toolset for whole-genome association and population-based linkage analysis. Am. J. Hum. Genet. 81, 559–575 (2007).

T. Jombart, adegenet: a R package for the multivariate analysis of genetic markers. Bioinformatics. 24, 1403–1405 (2008).

R Development Core Team, R: A language and environment for statistical computing (R Foundation for statistical Computing, Vienna, Austria, 2008).

S. Dray, A. Dufour, The ade4 package: implementing the duality diagram for ecologists. J. Stat. Softw. 22, 1–20 (2007).

E. Paradis, J. Claude, K. Strimmer, APE: analyses of phylogenetics and evolution in R language. Bioinformatics. 20, 289–290 (2004).

A. Raj, M. Stephens, J. K. Pritchard, FastSTRUCTURE: Variational inference of population structure in large SNP data sets. Genetics. 197, 573–589 (2014).

G. Evanno, S. Regnaut, J. Goudet, Detecting the number of clusters of individuals using the software STRUCTURE: a simulation study. Mol. Ecol. 14, 2611–2620 (2005).

R. J. Petit, L. Excoffier, Gene flow and species delimitation. Trends Ecol. Evol. 24, 386–93 (2009).

